# Functional Annotation Workflow for Fungal Transcriptomes

**DOI:** 10.64898/2026.01.12.697269

**Authors:** Nagisa Morihara, Hidemasa Bono

## Abstract

Although RNA sequencing (RNA-seq) enables rapid transcriptome profiling, functional annotation of fungal transcriptomes remains challenging. Existing tools prioritize broad taxonomic coverage, and reference genomes are scarce for non-model species. This study aimed to develop a fungal-specific functional annotation workflow to support rapid and accurate functional analyses downstream of RNA-seq, independent of reference genome availability. To evaluate the workflow, RNA-seq data from 57 samples of *Lentinula edodes* strain H600 (shiitake mushroom) were retrieved, along with full-length transcript sequencing (Iso-Seq) data and corresponding RNA-seq data from 20 samples of *Phakopsora pachyrhizi* (Asian soybean rust) from public databases. The workflow successfully annotated over 96% of protein-coding transcripts and demonstrated applicability to Iso-Seq data. Functional enrichment analyses revealed higher-resolution functional detection than existing annotation tools. Furthermore, integrating homology searches against fungal-specific databases with expression pattern-based annotations highlighted the workflow’s utility for target identification in genome editing and other applications. Overall, the results of this study highlight the potential of the developed workflow in facilitating the discovery of functionally important transcripts and their translation into biotechnological applications.

## 1. Introduction

Fungi represent one of nature’s most diverse organismal groups, with their exceptional functional diversity and ecological importance increasingly recognized in biotechnology, agriculture, environmental conservation, and human health [1]. Next-generation sequencing (NGS) technologies have enabled rapid, cost-effective genome sequencing, driving an exponential increase in sequenced fungal genomes. These resources offer opportunities to identify novel genes as targets for genome editing and functional characterization. RNA sequencing (RNA-seq), a key NGS application, is widely used to obtain transcript sequences and perform comparative transcriptome analyses across various conditions, time points, and treatments. This approach reveals actively expressed genes and their expression dynamics, providing direct insights into regulatory mechanisms and aiding research target prioritization. However, genome or transcriptome sequencing alone cannot elucidate gene function, as accurate functional annotation is essential for biological interpretation. Inaccurate or incomplete annotations hinder downstream analyses, such as functional enrichment studies and genome editing target identification [2]. Fungal functional annotation presents multiple inherent challenges compared with that in well-studied model organisms. First, many fungal species lack reference genomes, requiring reliance on phylogenetically distant or poorly annotated species; intraspecific diversity further necessitates comparative analyses across multiple strains. Second, fungi exhibit distinctive features, including biosynthetic gene clusters and species-specific adaptations, that are often absent in species-agnostic protein or domain databases, resulting in failure to capture. Finally, many predicted fungal genes lack experimental validation and uncharacterized with uncertain functional predictions. Consequently, annotation based on these resources often results in failure when analyzing closely related homologs lacking functional annotation [3, 4]. For these reasons, large-scale fungal sequence data have not yet been fully utilized.

Gene annotation involves identifying loci in genome sequences and assigning structural and functional information, including gene structure prediction, coding/non-coding region delineation, and functional inference based on sequence homology and existing functional data. Existing annotation workflows, such as BRAKER3 [5] and MAKER2 [6] for eukaryotic genomes, the Prokaryotic Genome Annotation Pipeline [7] for bacterial genomes, and funannotate [8] and FunGAP [9] for fungi, are often designed for submission to public repositories, including the National Center for Biotechnology Information (NCBI) [10], and therefore emphasize genome DNA-based structural prediction and gene determination. Meanwhile, several analytical tools are available for functional annotation of RNA-seq-assembled transcripts, including Trinotate [11], which provides multi-database annotation for model/non-model organisms; Blast2GO [12], which enables Gene Ontology [GO] enrichment *via* graphical user interface; and Fanflow4Insects [13], an insect-specific annotation tool. However, no dedicated tools exist for fungal transcriptomes.

In this study, a fungal functional annotation workflow applicable to transcriptomes with or without reference genomes was developed. By integrating fungus-optimized homology search databases and condition-specific annotations (e.g., tissue type and developmental stage), it aimed to address fungal-specific challenges, improving the quality and efficiency of downstream analyses, such as functional enrichment and genome editing target identification. Additionally, the applicability of this workflow was evaluated for full-length transcript sequencing (Iso-Seq) [14]. Iso-Seq enables high-resolution identification of splicing isoform structures, and data submissions to public repositories are expected to increase. The findings of this study demonstrate the workflow compatibility with both short-read RNA-seq and long-read transcripts, which highlight its versatility.

## 2. Materials and Methods

### 2.1 Acquisition of Expression Data

RNA-seq data for 57 samples of *Lentinula edodes* strain H600 were obtained from the Sequence Read Archive (SRA) (accession numbers SRR21185407–SRR21185463). Filtered, error-corrected Iso-Seq transcripts for *Phakopsora pachyrhizi* were retrieved from GenBank (accession numbers GHWK00000000 and GHWL00000000) [15]. Corresponding RNA-seq data, collected at days 3, 7, 10, and 14 post-infection, were obtained from the SRA (accession numbers SRR10130097–SRR10130116) [15].

### 2.2 Assembly and Coding Region Prediction

For the *L. edodes* dataset, raw reads from all 57 samples were merged and trimmed using TrimGalore (version 0.6.10) [16], followed by *de novo* transcriptome assembly with the rnaSPAdes module of SPAdes (version 3.15.5) [17]. Assembly quality was assessed using Benchmarking Universal Single-Copy Orthologs (version 5.8.0) [18], revealing 97.3% completeness. Coding regions were predicted using TransDecoder (version 5.7.1) [19]. For the *P. pachyrhizi* dataset, pre-assembled Iso-Seq transcripts were used directly, and short-read RNA-seq data were processed with TrimGalore.

### 2.3 Expression Quantification

Expression quantification was performed using Salmon (version 1.10.3) [20]. Treating each transcript as an independent unit, we used the ‘—keepDuplicates’ option during index creation to retain all variants. Transcripts Per Million (TPM) values were used for principal component analysis (PCA) in R (version 4.3.2) [21] with the stats and ggplot2 (version 3.5.2) [22] packages.

### 2.4 Functional Annotation

Workflow scripts are available in the fungifunate GitHub repository [23], built on the Systematic Analysis for Quantification of Everything framework [24]. Homology searches of retrieved coding region protein sequences used ggsearch36 from the FASTA package (version 36.3.8g) [25] with parameters ‘-d 1 -m10 -E 0.1’. Reference protein sequences for comparison were from human (*Homo sapiens*) [26], mouse (*Mus musculus*) [27], and budding yeast (*Saccharomyces cerevisiae*) [28], obtained from Ensembl [29], UniProtKB/Swiss-Prot [30], and FungiDB [31, 32]. Release68 protein files were concatenated into a single FASTA file, and protein domain searches followed using InterProScan (version 5.67-99.0) [33]. GO terms [34, 35] were retrieved for human, mouse, budding yeast, and UniProtKB protein IDs *via* the biomaRt package (version 2.58.2) [36] in R, and for InterProScan results using the InterPro2GO mapping file [37]. For the *L. edodes* dataset, condition-specific annotations were generated by averaging transcript-level TPM across the following three groups: mycelia (n = 19), primordia (n = 12), and fruiting bodies (n = 26). Stage-specific transcripts were defined as those expressed (mean TPM ≥ 1) in only one developmental stage, with low variability (coefficient of variation [CV] < 1) and maximum mean TPM ≥ These were consolidated into a single table using an R script.

### 2.5 Differential Expression Analysis

Differential gene expression analysis for *L. edodes* data used DESeq2 package (version 1.42.1) [38] in R, with pairwise comparisons across developmental stages. Wald tests identified differentially expressed transcripts, with statistical significance, using Benjamini– Hochberg adjusted *p*-values (padj) < 1 × 10^−11^.

For *P. pachyrhizi*, time-course expression patterns were analyzed with maSigPro package (version 1.74.0) [39] in R using third-degree polynomial regression. Significant genes (Benjamini–Hochberg adjusted Q-value < 0.05, ≥10 observations) underwent backward stepwise regression; those with R^2^ ≥ 0.6 were clustered into four groups *via* hierarchical clustering.

### 2.6 Functional Analysis

Functional enrichment analysis targeted differentially expressed transcripts using web tools Metascape (version 3.5) [40] and gProfiler [41] with human and budding yeast gene identifiers. Additionally, the topGO package (version 2.54.0) [42] in R analyzed all assigned GO terms and duplicates were removed. For topGO analysis, GO biological process (BP) ontology was examined *via* elim algorithms and Fisher’s exact test.

### 2.7 Comparative Annotation Method

Protein sequences were searched against the NCBI nr database [43] using DIAMOND (version 2.1.8.162) [44] in the blastp mode. GO terms were assigned with Blast2GO (version 6.0.3), followed by topGO functional enrichment analysis using the abovementioned parameters (section 2.6).

## 3. Results

### 3.1 Overview of the Functional Annotation Workflow

This workflow generated an annotation table from RNA-seq reads for functional enrichment analysis, integrating both functional annotations and differential expression results for transcript filtering. The functional annotation comprised four main components (Figure 1). First, homology-based annotation using a global alignment tool (ggsearch) against well-annotated protein databases, including human, mouse, budding yeast, UniProtKB/Swiss-Prot, and FungiDB. Second, GO term assignment for protein IDs from human, mouse, yeast, and UniProtKB. Third, protein domain annotation using InterProScan. Finally, condition-specific expression annotation.

**Figure 1.**
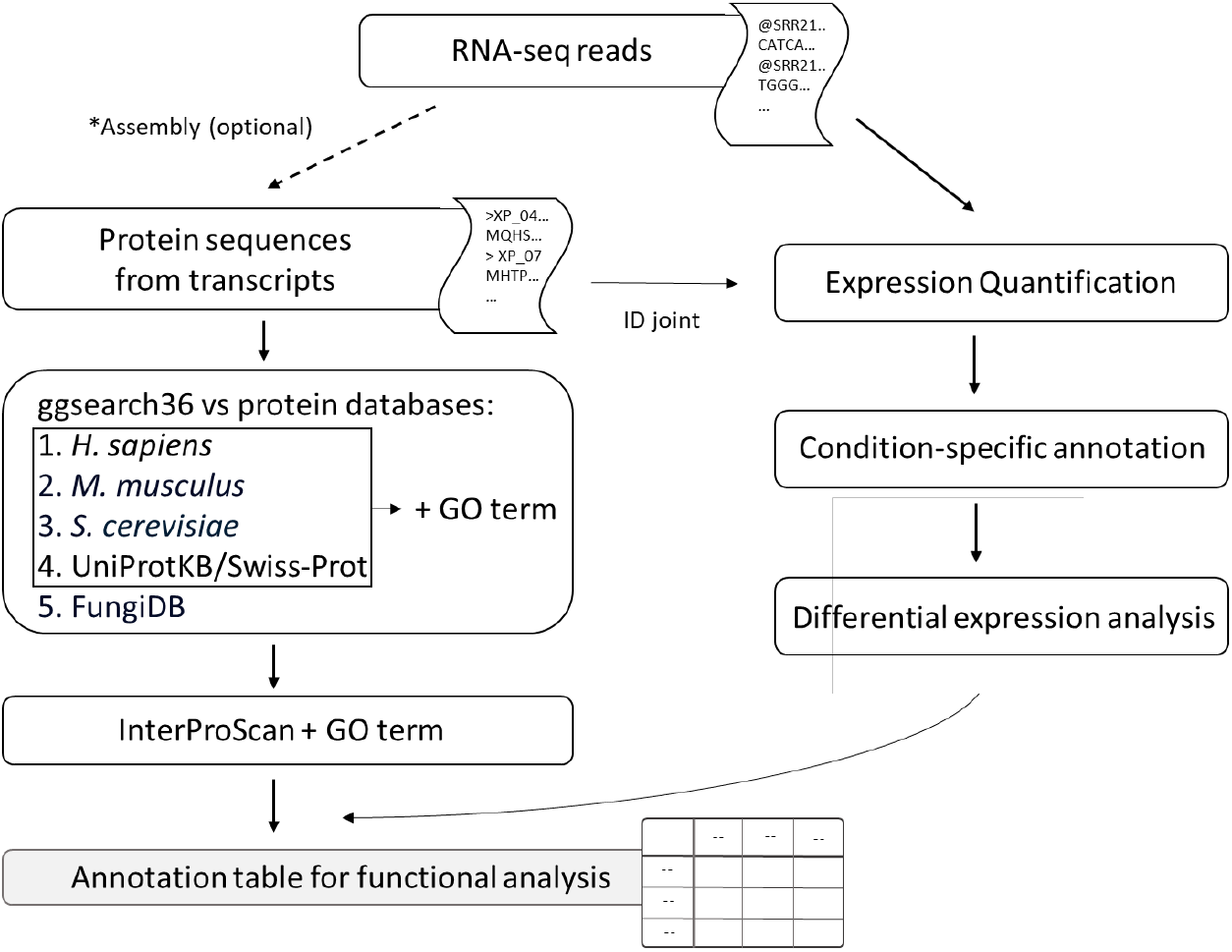
Overview of the annotation table generation including functional annotation. RNA-seq, RNA sequencing; GO, Gene Ontology.

Transcript sequences can be derived from *de novo* RNA-seq assembly and coding sequence prediction or from public databases. The ggsearch tool performs global alignments, making it suitable for detecting distant homologs in genetically diverse fungi GO terms provide a standardized framework for gene function across BPs, molecular functions, and cellular components, enabling functional enrichment analysis of significantly overrepresented functions. InterProScan identifies conserved protein domains and functional motifs, annotating transcripts missed by homology searches against protein databases and improving overall coverage. For condition-specific expression annotation, we applied custom binary criteria rather than continuous metrics, such as the Tau index [45], which are widely used to score tissue-specific expression on a 0–1 scale. A transcript was classified as condition-specific if it met all three criteria: (1) expression (TPM ≥ 1) in only one group, (2) CV < 1 across all groups, and (3) maximum mean TPM ≥ 2 in the expressing group. This binary TRUE/FALSE classification facilitates evaluation of expression patterns under specific biological conditions, such as developmental stages.

### 3.2 Application to L. edodes

To evaluate workflow utility, 57 RNA-seq samples from *L. edodes* strain H600 underwent *de novo* assembly and coding region prediction, followed by workflow application. These samples, originally classified into 20 groups by developmental stage and tissue type (Table S1) were consolidated into three broad groups (namely mycelia, primordia, and fruiting bodies) based on PCA (Figure S1) for condition-specific annotation and differential expression analysis. Annotation and expression data for each transcript were then integrated into a comprehensive table (Table S2). Of 227,580 analyzed transcripts, 98.2% received functional annotations. Contributions by database were as follows: human (42%), mouse (38%), budding yeast (30%), UniProtKB (55%), FungiDB (81%), InterPro (43%), condition-specific (0.13%), and GO terms (74%). In contrast, the NCBI nr database method annotated 66% of transcripts with only 12% assigned GO terms. Differential expression analysis identified 1,926 transcripts between mycelia and primordia, 1,739 between primordia and fruiting bodies, and 3,801 between fruiting bodies and mycelia. A Venn diagram (Figure 2a) was constructed, and functional enrichment analysis was performed for non-overlapping differentially expressed genes, including 492 specific to the mycelium versus primordium comparison and 711 specific to the primordium vs fruiting body comparison. Differential analysis between mycelia and primordia was performed through Metascape (Figure 2b) and g:Profiler (Figure 2c) analyses using human gene identifiers. This enabled the use of web-based tools that were previously inaccessible for non-model organisms. The results revealed significantly enriched metabolic processes (carboxylic acid, xenobiotic, and lipid metabolism) and functions related to oxidoreductase activity, cell membrane, and cytoskeleton. These results align with metabolic demands for primordium initiation and membrane/cell wall remodeling during mycelium-to-primordia transition [46-48]. Subsequently, GO enrichment analysis was performed using topGO with GO terms assigned by both approaches. As a result, the comparative method detected basic metabolic functions, such as carbohydrate metabolism and aromatic amino acid biosynthesis (Figure 2e), the other hand, our workflow detected more detailed functions related to cellular dynamics and specific pathways including NAD-cap decapping, fatty acid α-oxidation, microtubule-based peroxisome localization, and lamellipodium assembly regulation (Figure 2d). These differences reflect the differing GO term assignment rates between the two approaches.

**Figure 2.**
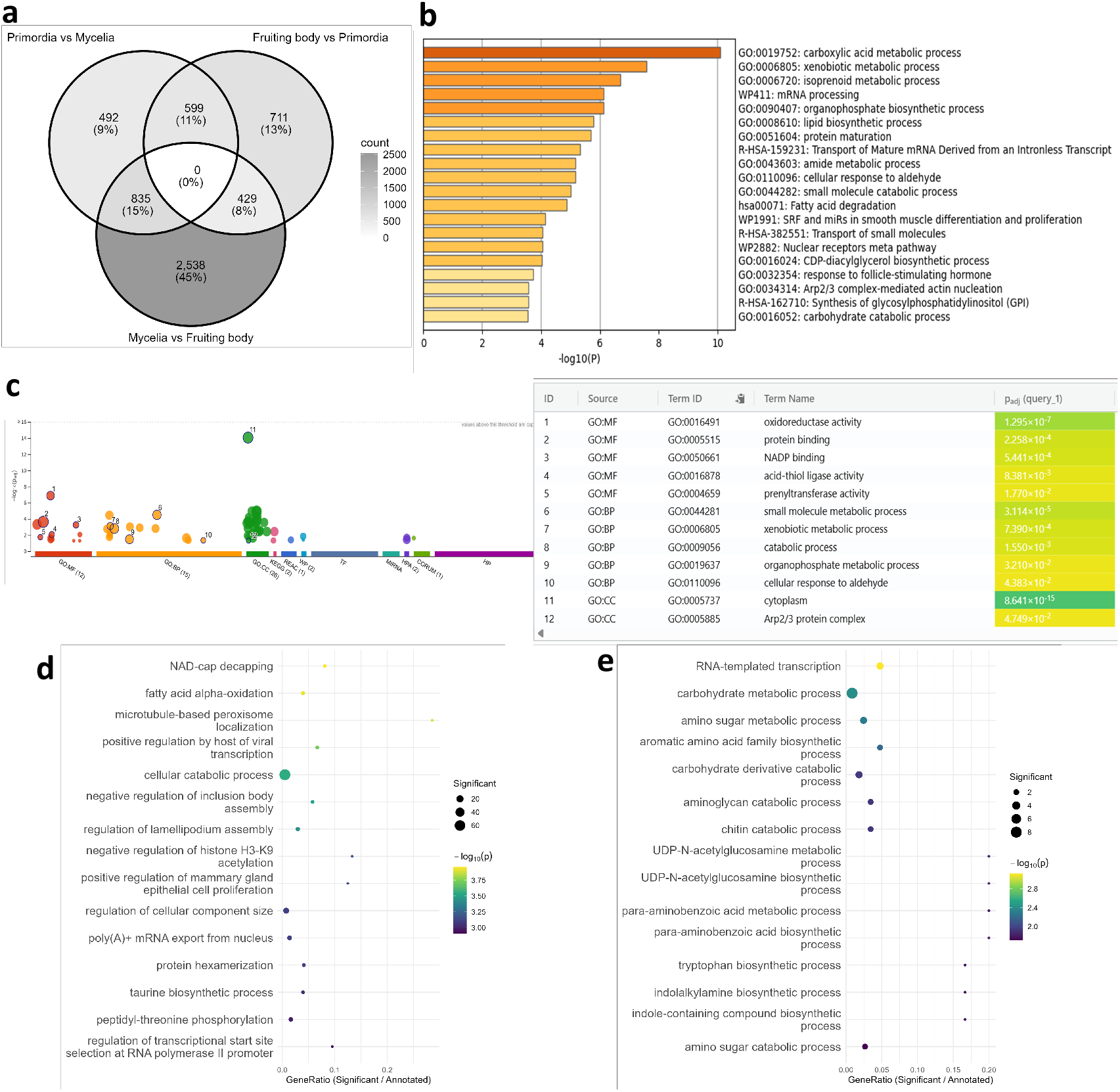
Functional enrichment analysis of *Lentinula edodes* transcriptome data. (a) Venn diagram of differentially expressed transcripts. The analyses shown in (b–e) were performed on the 492 transcripts uniquely differentially expressed between primordia and mycelia. (b) Metascape results using human gene identifiers. (c) g:Profiler results using human gene identifiers. (d) topGO dot plot based on GO terms aggregated from all annotation databases. (e) topGO dot plot based on GO terms assigned by the nr database-based method. GO, Gene Ontology; BP, biological process; CC, cellular component; MF, molecular function.

Between fruiting bodies and primordium, Metascape and g:Profiler analyses detected common enrichments in olefinic compound metabolism, xenobiotic metabolism, lysosomal lumen pH regulation, and iron ion-related functions (Figure S2a, b). Olefinic acid metabolism relates to unsaturated fatty acid metabolism, consistent with previously reported elevated expression during fruiting body development [49], along with potentially novel functions. Similarly, topGO analysis using our workflow identified mitosis, cell wall remodeling, and autophagy among top-ranked functions (Figure S2c), aligning with prior reports [50,51]. In contrast, the comparison method prioritized phospholipid biosynthesis, endoplasmic reticulum unfolded protein response, and microtubule-based nuclear migration (Figure S2d). Although these appear related to cell proliferation, they represented more ambiguous functional categories.

Notably, approximately 13% of transcripts were annotated exclusively *via* FungiDB, with many from uncharacterized basidiomycete genes, such as *Coprinopsis, Pleurotus*, and *Lentinus* (Table S2). Some of these appeared in differential expression analyses and considered priority candidates for genome-editing targets. Additionally, four transcripts received unique developmental stage-specific annotations. Among primordium-specific transcripts, a highly expressed homolog (TPM = 44) of yeast *SWS2*, involved in sporulation and oxidative stress responses, was identified (Table S2). This highlights the workflow’s ability to uncover functionally important genes beyond differential expression analysis.

### 3.3 Application to P. pachyrhizi

To evaluate workflow applicability to Iso-Seq data, filtered, error-corrected *P. pachyrhizi* Iso-Seq transcripts from GenBank were analyzed using the same pipeline. Annotation and expression data were consolidated into a single file (Table S3). Of 9,680 protein-coding transcripts, 96.1% received functional annotations, with contributions from each database as follows: human (56%), mouse (50%), budding yeast (44%), UniProtKB (55%), FungiDB (87%), InterPro (52%), and GO terms (79%). The comparative method annotated 80% of transcripts with only 19% GO terms assigned.

PCA confirmed separation of transcriptome data at days 3, 7, 10, and 14 post-infection (Figure S3). Time-course analysis identified 3,038 upregulated transcripts (Clusters 1–3) and 40 downregulated transcripts (Cluster 4) (Figure 3a). Functional enrichment was performed as described for *L. edodes*. For upregulated transcripts, both methods detected relatively broad functional categories (data not shown), which was attributed to their high proportion of upregulated transcripts compared with that in the total transcriptome. For downregulated transcripts, Metascape and g:Profiler identified CENP-A (histone H3 variant) and RNA-related processes (Figure 3b, c). Although histone modifications regulate pathogenicity [52] and host-derived histones exhibit antimicrobial activity [53], CENP-A associations with infection remain unreported. Subsequent topGO analysis showed that the comparative method prioritized nucleosome/ribosomal assembly and host-related functions (reductive pentose-phosphate cycle and photorespiration) as top enriched GO terms (Figure 3e). In contrast, the developed workflow detected specific, biologically relevant terms, including negative regulation of K48-linked ubiquitination, cell proliferation, and histone deacetylation (Figure 3d). These results confirm improved annotation accuracy for Iso-Seq data and applicability of the developed workflow.

**Figure 3.**
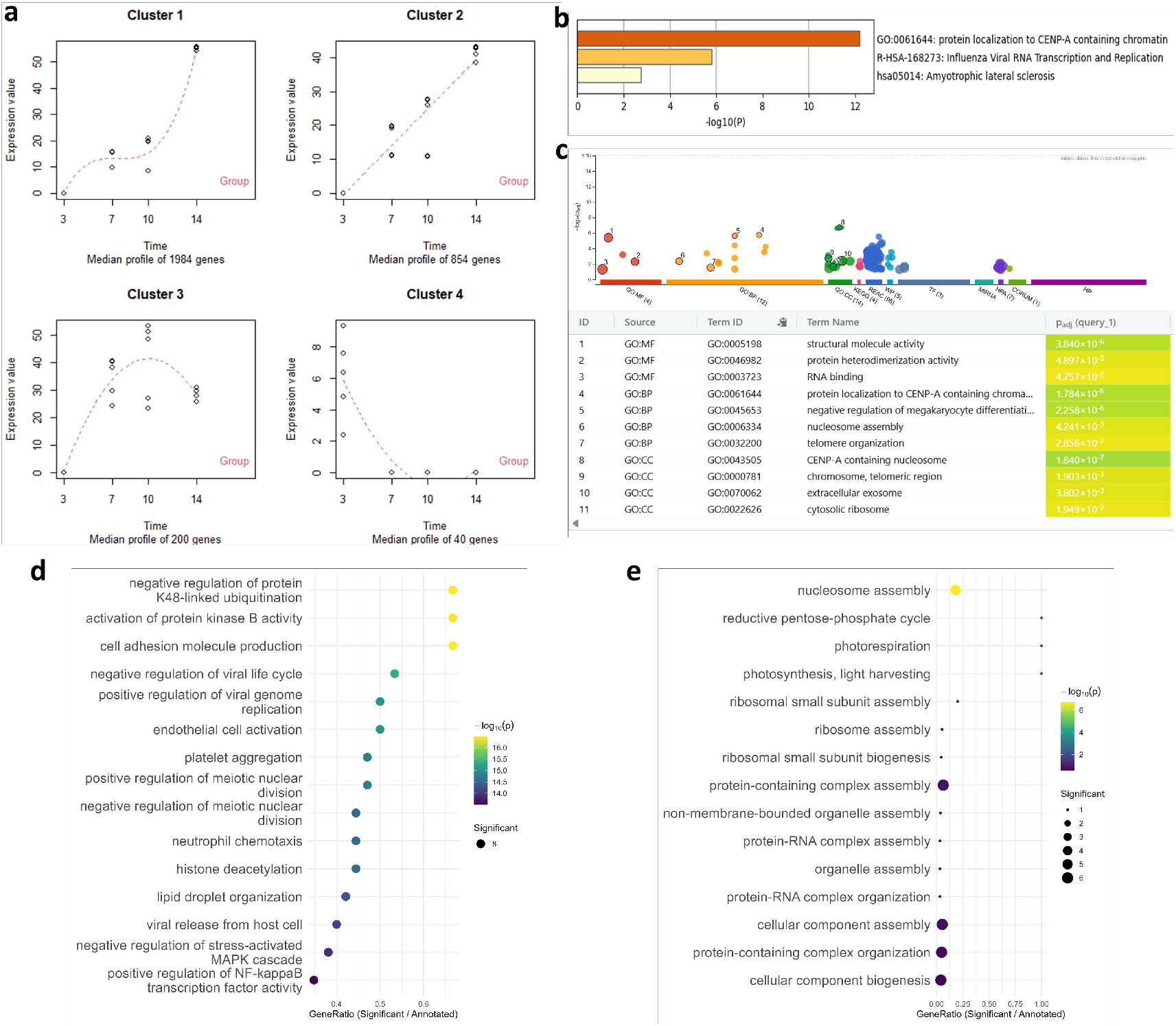
Functional enrichment analysis of *Phakopsora pachyrhizi* transcriptome data. (a) Cluster analysis of time-course differentially expressed transcripts. The analyses shown in (b–e) were performed on transcripts in Cluster 4. (b) Metascape results using human gene identifiers. (c) g:Profiler results using human gene identifiers. (d) topGO dot plot based on GO terms aggregated from all annotation databases. (e) topGO dot plot based on GO terms assigned by the nr database-based method. GO, Gene Ontology; BP, biological process; CC, cellular component; MF, molecular function.

Several differentially expressed transcripts received FungiDB-exclusive annotations, including uncharacterized genes from rust fungi and other pathogens (Table S3). These results suggest novel functions in rust fungi and represent future genome editing targets. Overall, the workflow delivers detailed, fungus-specific insights from both RNA-seq and Iso-Seq datasets.

## 4. Discussion

Publicly available RNA-seq data remain underutilized, particularly in fungi lacking comprehensive functional annotation. Reanalysis with the proposed workflow can uncover novel functional genes and response pathways. Even species with reference genomes may harbor unidentified splicing variants [54], making it valuable to utilize RNA-seq data-assembled transcript sequences. For *L. edodes*, the NCBI reference genome (GCF_021015755.1) contained 14,078 transcripts compared with our assembly result of 227,580 transcripts. Although redundant and non-translated sequences may exist, they may be filtered out during low-expression transcript removal or differential expression analysis. Moreover, the benefit of detecting novel transcript variants outweighs these limitations. In this regard, Iso-Seq analysis represents an effective approach. However, it should be noted that read number limitations may challenge complete coverage of all transcripts.

The developed workflow achieved superior annotation coverage and biological validity, particularly in GO term assignment. This was partially attributed to the use of NCBI nr database for BLAST results input to Blast2GO for comparison, which contains uncharacterized genes or entries lacking GO term associations. These results further emphasize the importance of incorporating databases with identifiers compatible with integrative functional analyses. Functional enrichment analysis using topGO revealed notable differences in biological insights between the two approaches. Differences originating from web-based tools, such as Metascape and topGO, may be partly attributed to the algorithm choice used in topGO. Herein, the elim algorithm was employed that preferentially detects more specific GO terms located at lower hierarchical levels. In contrast, Metascape and similar tools typically produce results closer to those generated by the classic algorithm, resulting in the enrichment of broader, higher-level parent terms. Additionally, web-based tools perform enrichment using gene identifiers from a specific model organism (e.g., human), whereas topGO integrates GO terms aggregated from all databases, further generating differences in the outputs. Results obtained for *L. edodes* and *P. pachyrhizi* showed that the proposed workflow identified more specific and narrowly defined functions, which were directly linked to core cellular dynamics, including transcript remodeling, the G2/M transition of the mitotic cell cycle, and ubiquitination, thereby efficiently annotating candidate genes for downstream functional validation. The workflow also detected lower-level GO categories involving relatively small numbers of genes; for example, the NAD-cap decapping category contained 37 genes, and the negative regulation of protein K48-linked ubiquitination category contained 12 genes, representing a practically manageable number of candidates for gene-focused analyses. In the topGO results for *P. pachyrhizi*, the detected broad host-associated functions included photorespiration and photosynthesis by nr-based annotation approaches, requiring careful interpretation. Although host derived sequences were removed at the transcript level, these annotations may arise from assignments to plant-related genes in the nr database or artifacts inherent to homology-based annotation. In contrast, the developed workflow utilized databases with explicit taxonomic constraints, including fungal-specific resources, model organism datasets, and UniProt, thereby reducing such artifacts. Notably, integrating GO terms from multiple databases reduces annotation bias, increases the detection of smaller gene-set categories, and supports more accurate biological interpretation. FungiDB contributed the most to annotation efficiency owing to its continuous integration of published and unpublished fungal data, including functionally unknown genes. This further suggests that several fungal genes remain undercharacterized and unregistered in species-agnostic databases. Annotations of differentially expressed transcripts exclusively by FungiDB were largely derived from closely related species, suggesting that FungiDB complements important fungus-specific information not captured by conventional annotation approaches. Developmental stage-specific annotation in *L. edodes* exclusively annotated four transcripts through this method. A high-expression homolog of a yeast oxidative stress response gene was detected among primordium-specific transcripts; however, it was excluded from the differentially expressed transcript list because of the applied q-value cutoff. This approach allowed prioritization of potentially important genes independent of other annotation strategies and differential expression analyses. Despite the low developmental stage-specific annotation rate (0.13%), analysis based on the original 20 classification categories may reveal additional stage-or tissue-specific transcripts.

The workflow developed in this study relies on homology-based annotation, making it dependent on reference database quality and imperfect cross-species functional inference. This underscores the need for experimental validation. Despite these limitations, opportunities exist for optimizing annotation strategies, refining database selection, and expanding the analysis. In a multi-database integration strategy, annotation order across these databases is important, as prioritizing well-annotated species may yield more useful functional information. This observed higher annotation rate from human sequences compared with those from phylogenetically closer yeast species may be attributable to the more extensive characterization of human transcript variants. Although this workflow integrates GO terms for facilitating easy functional interpretation and widespread use, its flexibility allows for the incorporation of additional database information per requirements, such as enzyme commission numbers, Clusters of Orthologous Groups/Eukaryotic Clusters of Orthologous Groups classifications, or secondary metabolite biosynthetic gene cluster annotations from resources, such as antiSMASH database. Although focused on protein-coding transcripts, expanding to non-coding RNAs (ncRNAs), such as long ncRNAs and micro RNAs, as key regulators and genome editing targets [55] would benefit from rRNA depletion-based library preparation, instead of mRNA-seq. Future directions of this study include machine learning-based structural annotation for both coding and non-coding transcripts, which is particularly valuable considering fungi’s high genetic diversity and low sequence homology.

In conclusion, the proposed workflow provides a practical, reference-genome-independent framework for fungal transcriptome functional annotation. It enables mechanistic investigation of underexplored, complex biological processes underlying developmental programs and environmental responses even in fungal species, capturing genes and functions missed by conventional approaches. Furthermore, prioritized candidates serve as rational CRISPR-based functional validation, facilitating the translation of annotated transcripts from basic to applied research.

## Supplementary Materials

The following supporting information can be downloaded from: https://doi.org/10.6084/m9.figshare.30944468, Figure S1: PCA plot of *L. edodes* RNA-seq datasets; Figure S2: Functional enrichment analysis of 711 uniquely differentially expressed transcripts between fruiting body and primordia in *L. edodes*; Figure S3: PCA plot of *P. pachyrhizi* RNA-seq datasets; Table S1: *L. edodes* dataset metadata; Table S2: *L. edodes* annotation table; Table S3: *P. pachyrhizi* annotation table.

## Author Contributions

Conceptualization, N.M. and H.B.; methodology, N.M. and H.B.; software, N.M. and H.B.; validation, N.M..; formal analysis, N.M.; investigation, N.M.; resources, H.B.; data curation, N.M.; writing—original draft preparation, N.M.; writing—review and editing, N.M. and H.B.; visualization, N.M.; supervision, H.B.; project administration, H.B.; funding acquisition, H.B. All authors have read and agreed to the published version of the manuscript.

## Funding

This study was supported by the Center of Innovation for Bio-Digital Transformation (BioDX), an open innovation platform for industry-academia co-creation (COI-NEXT), and the Japan Science and Technology Agency (JST), grant number JPMJPF2010.

## Institutional Review Board Statement

Not applicable.

## Informed Consent Statement

Not applicable.

## Data Availability Statement

The original data presented in this study are openly available on GitHub [23].

## Acknowledgments

Experiments were performed using computers at the Hiroshima University Genome Editing Innovation Center.

## Conflicts of Interest

The authors declare no conflict of interest. The funders had no role in the design of the study; in the collection, analyses, or interpretation of data; in the writing of the manuscript; or in the decision to publish the results.

## Abbreviations

The following abbreviations are used in this manuscript:

TPM: Transcripts Per Million
RNA-seq: RNA Sequencing
Iso-Seq: Full-Length Transcript Sequencing
NGS: Next-Generation Sequencing
NCBI: National Center for Biotechnology Information
SRA: Sequence Read Archive
PCA: Principal Component Analysis
CV: Coefficient of Variation

## Notes

https://doi.org/10.6084/m9.figshare.30944468

### Competing Interest Statement

The authors have declared no competing interest.

### Summary of Updates

Some Figure citations were missing in the previous version, and those are fixed in this version.

